# Model-based inference of enzyme inhibitions from perturbation-induced metabolic dynamics

**DOI:** 10.64898/2026.06.01.729220

**Authors:** Wolfram Liebermeister, Michela Pauletti, Terézia Dorčáková, Caspar Rahm, Mattia Zampieri

## Abstract

In microbes, metabolism plays a key role in the first rapid adaptation to sudden challenges such as nutrient limitations or toxic compounds. While metabolomics enables the profiling of stimuli-induced metabolic changes, computational approaches that can interpret these data to mechanistically explain how perturbations propagate through metabolism to produce the observed changes are lagging behind. Here, we developed a computational framework, called Inference from Metabolic Fingerprints (IMF), to model the immediate dynamic response to a metabolic perturbation and systematically infer its entry point (i.e. enzymatic target). IMF assumes small perturbations and linearizes the nonlinear dynamics around a reference steady state. This allows IMF to scale with large metabolic networks and bypass missing kinetic parameters by allowing for fast and efficient ensemble sampling. We apply IMF to a model of central metabolism in *Escherichia coli*. Using in-silico and experimental data, we demonstrate the ability to infer the target of metabolic perturbations in spite of unknown kinetic parameters and incomplete metabolic data. Hence, we show that IMF is an effective approach for designing, analyzing and interpreting time-resolved metabolomics.

## Introduction

Cells respond to environmental challenges such as stress, changes in nutrient composition or toxic compounds in a variety of ways and on different time scales. Bacteria surviving extreme low temperature, high pressure or rapid changes in pH and redox state, or cancer cells extravasating, surviving immune response and colonizing a different tissue are emblematic examples of cells phenotypic plasticity – i.e. the ability of the same genotype to show different phenotypes as a function of the environment ^1–4^. While successful long-term adaptation largely depends on changes in gene expression, mounting evidence suggests that immediate metabolic response is crucial in mediating adaptation and thereby cell fate ^5–7^.

Extrinsic perturbations such as drug treatments will cause a pervasive metabolic “echo” that propagates from a directly affected target (e.g. an enzyme) to more distal parts of metabolism^8–10^. Enzyme inhibitors will induce metabolic changes that go beyond the simple accumulation of substrates and depletion of products in the perturbed reaction. Instead, these immediate changes are rapidly transmitted to neighboring or distal reactions by shifting reactions equilibria or by changing kinetic properties of enzymes through allosteric regulation. To understand how metabolism responds to an external perturbation (e.g. drug), one needs to systematically describe the interplay between the entry point of the perturbation (i.e. drug target) and its interactions within the systems network (kinetic and regulatory).

New high-throughput mass spectrometry technologies enable the profiling of metabolic changes at a time resolution of seconds and with a high coverage of measured metabolites ^11–16^. Metabolic profiling is routinely used to monitor dynamic changes in the abundance of hundreds of metabolites in response to genetic or external (e.g. drug) perturbations ^8,9,17–20^. The bottleneck has entirely moved from data generation to the need for predictive models able to extract mechanistic insights from large metabolomics data sets. Distinguishing direct metabolic consequences of a perturbation, such as those produced by the direct interaction of a drug with the target, from indirect changes, such as those induced by adaptive cell regulatory mechanisms, is of utmost importance to shed mechanistic insights on the mechanism of action of perturbations, such as drugs, and predict possible side effects or optimal drug combinations ^10,21–29^. However, identifying the direct target(s) of a perturbing agent directly from perturbation-induced metabolic changes remains challenging.

At first sight, inferring inhibited enzymes from their “metabolic echo” may seem easy. For linear, unregulated pathways, the crossover theorem ^30–32^ predicts that metabolites upstream of an inhibited enzyme will accumulate and metabolites downstream will deplete. Under ideal conditions, by simply displaying the measured metabolite fold changes on a metabolic network, we should be able to identify the inhibited enzyme by visual inspection. However, in practice things are more complicated: metabolic networks contain cycles, and seemingly distant reactions may be coupled via cofactors or regulation of enzymes. Moreover, metabolomics measurements are noisy and cannot resolve all metabolites in the network. The question is whether under such real-life conditions - and even when a perturbed reaction is not immediately visible from the data - the entire metabolite profile, measured shortly after a perturbation, contains enough information to infer the source of the perturbation. In this context, we refer to the enzyme directly affected by the perturbing agent as the target, whereas the resulting change in enzyme activity is the source of the downstream perturbation. Kinetic metabolic models provide a tool to simulate perturbation-induced metabolic changes and to analyze dynamic metabolome data^33–36^. In principle, by fitting a model to experimental data recording metabolic changes in response to an external perturbing agent, one might reveal the direct target of the perturbation and the details of its propagation through the metabolic network. However, a large computational effort and large numbers of unknown parameters limit the scope of kinetic models to model organisms and small-scale networks^37–44^.

How can we scalably simulate the dynamic response of metabolite concentrations to perturbations? Given the flux directions and magnitudes in an unperturbed metabolic state, the dynamic metabolic response to an enzyme perturbation depends crucially on how strongly perturbations are transmitted from one reaction to its downstream and upstream neighboring reactions. This depends on the reaction elasticities, which quantify how strongly a reaction rate is directly impacted by concentration changes in the reaction or by concentration changes of effector molecules regulating the enzyme. On a larger scale, the propagation of perturbations is largely determined by structural facts such as network structure, regulation arrows, and flux directions.

In signaling networks, observed temporal dynamics have been used to infer the temporal shape of the original perturbation along with the set of model parameters ^45^. Here we tackle a similar problem, but instead of determining the shape of the perturbation at a known input node, we assume a step-wise perturbation function and ask which of the nodes has been perturbed. At the same time, we take inspiration from works on signaling and metabolic network that have used notions from Metabolic Control Analysis to estimate reaction elasticities - and therefore, pairwise interactions between network nodes - from the steady-state effects of perturbations ^46,47^. However, instead of considering steady-state changes, we consider temporal changes described by dynamic control coefficients, and instead of estimating the elasticities, we treat them as nuisance parameters to be sampled, using the Structural Kinetic Modeling (SKM) framework^48–50^.

SKM provides a direct way to study the roles of local kinetics and global network structure in metabolic dynamics. The basic idea is to define a model based on structural knowledge and to explore its properties by repeatedly sampling unknown parameters from random distributions. Reparametrizing the model with elasticities instead of kinetic constants greatly simplifies model construction and simulation and allows for an efficient exploration of the kinetic and regulatory parameters space, taking various types of information into account. While the precise values of reaction elasticities may be largely unknown, educated guesses or random sampling can be used to simulate the most plausible ways in which perturbations propagate in a network. Comparing these expected dynamic patterns, or “metabolic fingerprints”, to measured metabolite data can help us infer the most probable causes of observed, dynamic perturbations.

Using this method, we developed a novel framework called Inference from Metabolic Fingerprints (IMF) to systematically infer enzymatic targets of a perturbation from dynamic metabolome profiling data. While classical kinetic models employ systems of nonlinear differential equations, IMF linearizes the dynamics around the initial unperturbed state, which makes the method versatile and scalable. Below, we first describe the mathematical formulation of IMF, the assumptions made, and the information required to simulate dynamic metabolic response to perturbations. We then test IMF on simulated perturbations on a model of central carbon metabolism in *E. coli* and benchmark its predictive power by testing sensitivity to noise and coverage (i.e. the number of measured metabolites). Finally, we test our approach on experimental mass spectrometry data monitoring the response of *E. coli* to hydrogen peroxide and demonstrate its ability to infer the enzyme inhibitions that best explain the observed immediate response to oxidative stress.

Matlab code for IMF is available on GitLab (see Code Availability).

## Results

To infer perturbed enzymes from their immediate metabolic responses - in the first few minutes - we developed a modeling framework that simulates metabolic dynamics in a linear approximation. In the inference procedure, we compare measured perturbation-induced metabolic changes to the changes predicted by simulations. We then assign to each enzyme a probability of being the source of the perturbation, together with an estimate of the inhibitory strength. The probabilities account for uncertainties both in metabolomics data and in model parameters. After testing with artificial data, we applied IMF to metabolic dynamics observed in an oxidative stress experiment in *E. coli* bacteria ^6^.

### Model construction, simulation, and inference of perturbed enzymes

To simulate metabolic dynamics after an enzyme perturbation, we use kinetic models based on Ordinary Differential Equations (ODEs). In classical kinetic models, the reactions are described by nonlinear rate equations^51^ such as Michaelis-Menten rate laws or modular rate laws^52^, which can also account for small-molecule regulation of enzymes^14,53^. Since such models would require time-consuming numerical integration, we simplify the nonlinear equation system by a linear approximation around the initial, unperturbed state (see Methods and Supplementary Figure 1). Following earlier work on temporal metabolic responses^54^, we then describe the metabolic response to a sudden shift in enzyme activity by time-dependent control coefficients, which can be efficiently computed by matrix exponentials. For long times (i.e. t->∞) after the perturbation, the time-dependent control coefficients converge to the static control coefficients used in Metabolic Control Analysis^55^.

Compared to classical kinetic modeling with explicit numerical integration, the proposed approach allows not only for faster calculations, but also for an efficient sampling of the space of possible model parameters. The resulting model ensemble provides us with uncertainty ranges for all model results, while all model instances share the same network structure and the same unperturbed metabolic states.

By simulating different enzyme inhibitions in a model and maximizing the similarity between these simulations and experimental data, we can generate experimentally testable hypotheses about the target of a perturbation into metabolism. Hence, the proposed approach opens the door for new and effective ways to understand, mechanistically, how toxic compounds or drugs can directly affect metabolism and what are the regulatory mechanisms mediating the immediate response, based on metabolic profiling of perturbation-induced metabolic changes.

To generate an approximated linear model of cell metabolism and simulate its response to external perturbations we necessitate a stoichiometric model and an initial metabolic steady state obtained from experimental or computational inferred absolute measurements of metabolites and fluxes. There exist various manually curated stoichiometric models for bacteria, plant and animal cells^56–58^ as well as tools to generate stoichiometric models directly from genome sequences^59^. Although information about direct enzyme regulation^14,53,60^ (e.g. allosteric regulation or competitive inhibition) can be directly used in our computational framework, in the following we benchmark IMF using only the stoichiometric structure of the metabolic network, that is, assuming that reaction rates depend only on substrate, product, and enzyme concentrations.

Absolute metabolite concentrations and fluxes remain difficult to obtain experimentally. Estimating intracellular fluxes typically requires the use of labelled substrates and is restricted to chemically defined conditions with unique carbon and nitrogen sources^61,62^. Moreover, measurement values of fluxes and metabolite concentrations will be sparse - i.e. not all fluxes or metabolites can be experimentally measured. However, using constraint-based modeling and thermodynamic constraints, gaps in the data can be filled^38,63^. Here we do this by projecting fluxes to a steady-state flux distribution in our model and choosing complete and thermodynamically consistent metabolite concentrations by using a variant of parameter balancing^64^ .

Once a reference state has been determined and reaction elasticities have been chosen, the model allows us to simulate the effect of virtually interfering with any reaction(s). In the following, we model enzyme perturbations by applying partial (percentage) inhibitions to individual enzymes.

If the reaction elasticities of our model were already known, simulating the effect of a perturbation would be straightforward. Here, we assume that they are completely unknown and treat them as nuisance parameters. By sampling the reaction elasticities, we can generate a large ensemble of model instances and study their dynamics in response to an enzyme inhibition. Elasticities can be sampled (or chosen) based on an independent sampling (or choice) of the underlying saturation values^50^. The formula for converting saturation values into elasticities depends on the assumed underlying enzymatic rate laws, and in the case of reversible rate laws, the elasticities additionally depend on the thermodynamic driving forces. Each choice of saturation values yields a complete full set of kinetic parameters, and hence simulations can either be run with fully parametrized non-linear kinetic models or directly with the linearized model defined by the elasticities.

The dynamic response of metabolite concentrations to a perturbation is described by a “metabolic fingerprint” - a profile of simulated or measured metabolite log-concentrations, for a set of metabolites and a number of time points. A metabolic fingerprint, as a data matrix, can be visualized as a heatmap (see Figure 1C). By comparing simulated and measured metabolite fingerprints, we can infer the perturbed enzyme and set of elasticities that most accurately describe the experimental data.

**Fig. 1.**
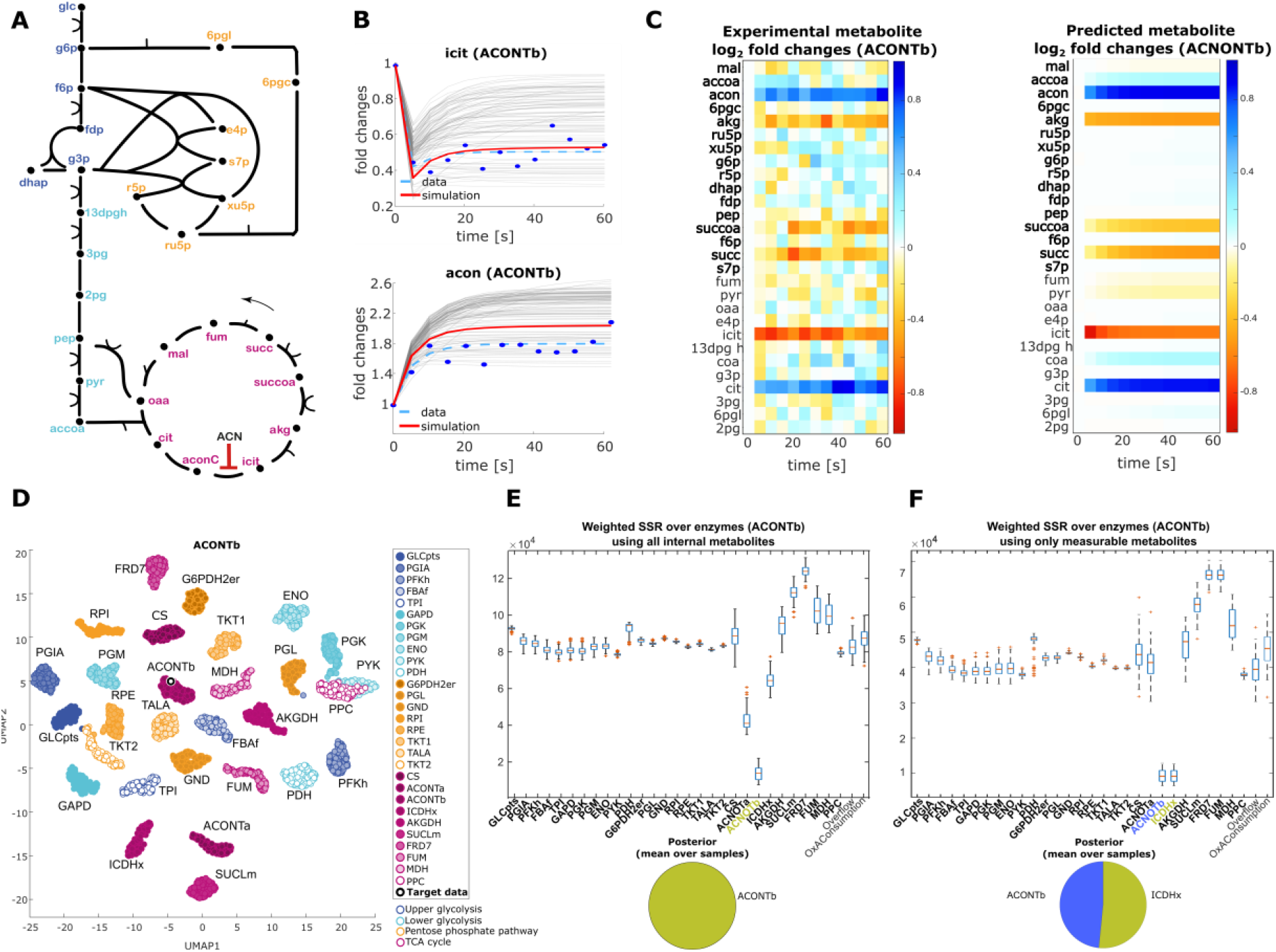
Simulation of metabolic dynamics and inference of perturbed enzymes. To simulate the dynamics of all the metabolites in 13 time points (0 - 60 seconds) after the onset of the inhibition, we applied a 50% inhibition to aconitase (ACN) and constructed the target data by adding gaussian noise on logarithmic scale (geometric standard deviation of 1.1., corresponding to a noise level of approximately 10 %). The enzyme kinetics were described by reversible Michaelis-Menten type of equations with predefined kinetic parameters (i.e. K_M_ and V^max^)). Next, we generated a hundred model instances with sampled reaction elasticities, and simulated for each of them the inhibition of each enzyme in the network. The weighted sum of squared residuals (wSSR) between all the metabolite dynamics in the target data and in each simulation was calculated and the best matching reaction was identified. **A** - The *E. coli* model of metabolism used to generate the data contains 31 reactions and 40 metabolites (out of which 12 are considered external, modeled with fixed concentrations). **B** - The dynamics of the product (isocitrate - icit) and the substrate (cis-aconitate - acon) of the inhibited ACN reaction. Grey lines represent the icit/acon dynamics in all simulations of ACN inhibition, the red line highlights the simulation which best matches the target data, the light blue dashed line represents the target data, and the blue dots indicate the target data dynamics after adding noise. **C** - The heatmaps show the dynamics of all the internal metabolites of the target dataset (left) and the best matching simulation (right). **D** - UMAP projection of the metabolic fingerprints from all hundred model instances, considering inhibitions of each reaction in the network. Each dot represents one metabolic fingerprint, and nearby dots correspond to similar fingerprints. Fingerprints from the target data are shown by circles. **E, F** - Each of the fingerprints is characterized by a log-likelihood value (the negative sum of squared residuals), describing its similarity to the target data. The box plots show distributions of the negative log-likelihoods, where each box refers to a perturbed enzyme tested. The log-likehoods were computed either using all the internal metabolites (E) or using only metabolites assumed to be measurable (measurability determined by the dataset from figure 6, highlighted in the heatmaps (C) in bold) (F). The pie charts show the posterior probability of each enzyme to be the source of the perturbation. Only enzymes with the smallest negative log-likelihood values obtain substantial posterior probabilities. For more details, see Supplementary Figures.

In principle, we could simulate the metabolic fingerprint of each enzyme, one by one, in the model, and choose the one that best resembles the measured fingerprint. The similarity between fingerprints could be assessed by distance measures such as the Euclidean distance. However, to properly take uncertainties into account, missing data, and potentially prior knowledge, we treat the problem as a statistical inference problem. Using the measured data, we obtain approximations of the posterior probability for each enzyme to be responsible for the observed metabolic changes. Instead of running a full Bayesian inference procedure with posterior sampling, we use a simpler and faster algorithm based on samples from a prior distribution (see Methods). Based on the posterior probabilities, our algorithm then proposes a single enzyme or a number of most plausible enzymes to be the target of the perturbation.

### In silico benchmarking of drug target IMF

To test IMF and to quantify its accuracy, we first used in-silico data generated from a sudden inhibition of an individual enzyme in a model of *E. coli* central metabolism, but with full kinetic rate laws. The model is a modified version of the model from ^65^ that consists of 40 metabolites and 31 reactions (Figure 1A). To define an initial, unperturbed reference state, we constructed a thermodynamically consistent steady state based on experimentally determined fluxes^66^ and metabolite concentrations^67^ in *E. coli* bacteria exponentially growing in a glucose minimal medium. Since only some of the metabolites and fluxes can be measured experimentally, we adopted a computational strategy to infer missing flux measurements by using flux balance analysis, and to infer missing metabolite concentrations by using parameter balancing^68^ to determine concentrations providing feasible Gibbs free energies in all the reactions.

Using this unperturbed reference state, we simulated metabolite dynamics based on a model with reversible Michaelis-Menten rate laws and a fixed set of kinetic parameters (i.e. K_m_ and V_max_) obtained via elasticity sampling. We simulated a 50% inhibition of enzyme activity (i.e. step function on a single enzyme) and computed the metabolite profiles at 13 time points up to 60 seconds after the perturbation (0-60 secs). Independent Gaussian noise was added on logarithmic scale to simulate measurement errors (Figure 1B-C and Supplementary Figure 2).

Next, we used the 31 time resolved, in silico generated metabolic fingerprints to benchmark IMF and to assess its ability to infer the inhibited enzyme. To this end, we computed a posterior probability for each enzyme in the model, indicating how plausible it is that this was the enzyme perturbed. An enzyme’s posterior depends on how similar its simulated fingerprint is to the “true” metabolite fingerprint and, possibly, on prior probabilities for the different enzymes (for simplicity we used a uniform prior), and captures all uncertainties in our model. The uncertainties in our model arise not only from simulated measurement errors, but also from the fact that the true reaction elasticities are unknown. In a Bayesian setting, these elasticities could either be treated as variables to be estimated or as nuisance variables. Here we account for them as nuisance variables and account for them simply by sampling.

To do so, we sample elasticities, giving rise to an ensemble of 100 model instances with different Jacobian matrices and corresponding kinetic parameter sets, consistent with the respective steady states. It is worth noting that model instances with an unstable reference state were discarded. For each of the 100 model instances leading to stable states, we simulated the systems’ dynamic responses to the inhibition of each individual enzyme (Figure 1B-C). Specifically, we use the linearized model and 100 sampled Jacobians (each obtained from a sampled scaled elasticity matrix) to simulate dynamic metabolic changes upon inhibition of enzyme activity (Figure 2C). Based on the simulation runs, we estimate the likelihood of each enzyme, in each model instance, to be the target of the perturbation (Figure 1D). The log-likelihood results from a weighted Euclidean distance between the simulated and “true” metabolite fingerprint. We then aggregate the results for the 100 model instances into a single likelihood for each enzyme. Assuming that all enzymes are equally likely to be perturbed a priori, this likelihood yields the posterior, stating how plausible it is that this was the enzyme perturbed.

**Fig. 2.**
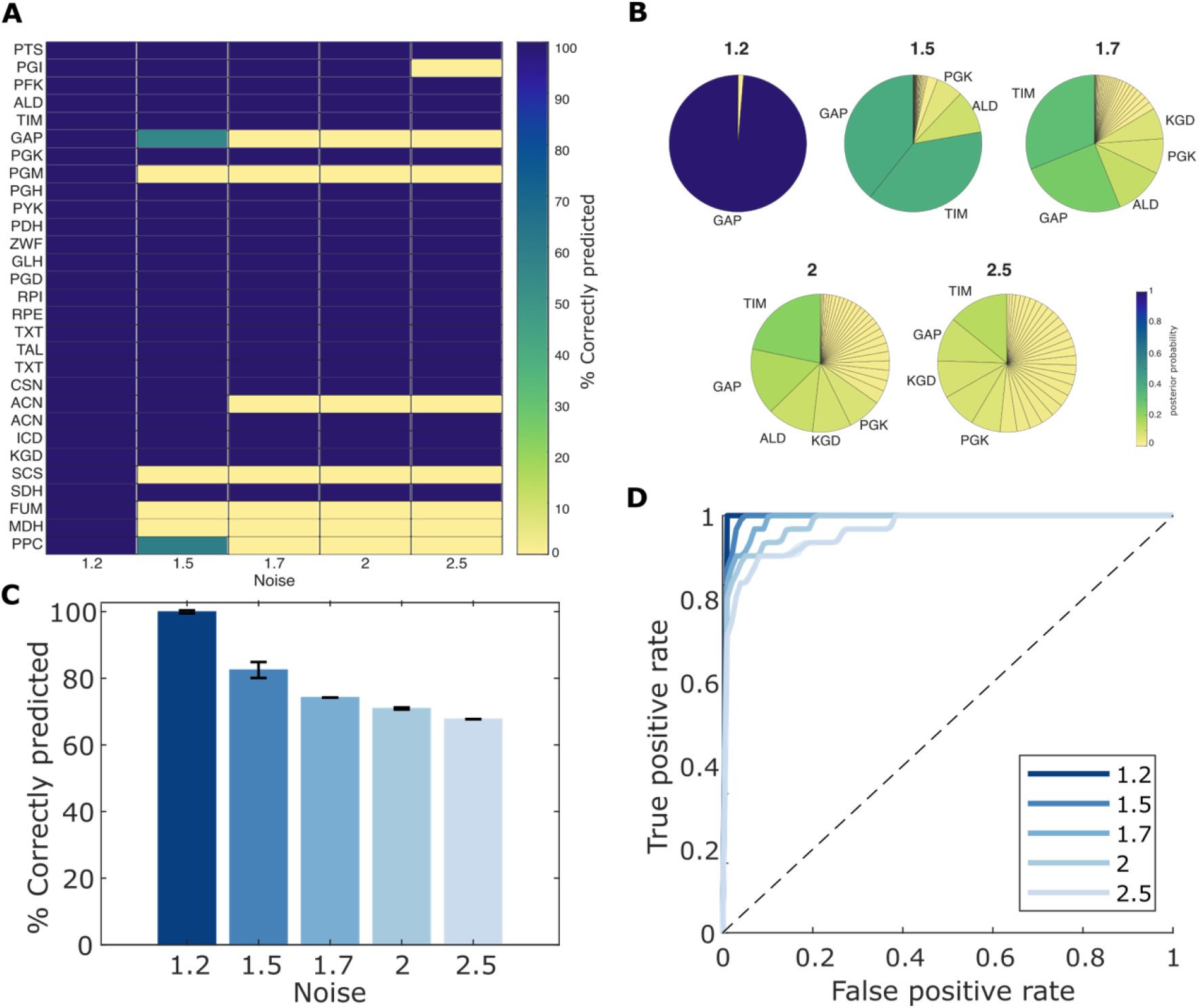
Statistics on the inference of inhibited reactions with artificial data, simulating inhibitions of each of the enzymes. As previously (figure 1), we simulated dynamics of all metabolites in 13 time points within 0-60 seconds following a 50% inhibition of each reaction at a time. We inferred the inhibition of each reaction in the network a hundred times for five different noise levels. **A** - The heatmap displays the percentage of correct predictions across individual reaction and noise levels, revealing that most reactions remain robust to noise, while the performance rapidly drops under higher noise only for a small subset. **B** - Pie charts indicate posterior probabilities of reactions when inferring GAP inhibition given different noise levels. Only reactions with probability > 0.05 are labeled. **C** - The bar plots represent the percentage of correctly inferred perturbed reactions given each noise level. The prediction accuracy consistently reached 100% at the lowest noise level (with a geometric standard deviation 1.2, corresponding to about 20%) and declined as noise increased. **D** - The receiver operating characteristic (ROC) curves indicating the prediction accuracy across different noise levels.

To benchmark our inference method, we assessed how often the enzyme predicted by our model - that is, the one with the highest posterior probability - matched the true inhibited enzyme – i.e. the enzyme inhibited to generate the true in silico dynamics (Figure 2A-B). Aside from only checking the enzyme with the highest posterior, we also asked if the true perturbed enzyme appears in the first *n* top-level predictions based on the posterior probabilities of all the enzymes. To assess this, we performed a ROC analysis comparing in silico perturbed enzymes with model predictions, estimating false and true positive rates (Figure 2C-D). In ROC analysis, curves that rise steeply toward the upper-left corner indicate better discriminative performance, with the area under the curve (AUC) summarizing overall accuracy. To examine how inference performance degrades with increasing measurement uncertainty, we repeated the analysis across a series of simulated increasing noise levels. Consistent with expectations, increasing the noise levels or reducing the number of detectable metabolites - in this case: metabolites used for comparing in silico data to IMF simulations - led to a substantial decline in prediction accuracy (Figure 2).

Overall, we found that for a noise level around 1.2. (corresponding to a measurement error of about 20%, similar to actual average noise levels in time-resolved metabolomics data ^69^, IMF exhibits remarkably high accuracy (i.e. AUC=1). Specifically for all 31 simulated enzyme inhibitions IFM was able to infer the correct perturbation target. In some cases, the ability of IMF to infer the target of the perturbation was supported by clear local effects: strong accumulation of upstream enzyme substrates and depletion of downstream products. For example, simulating the effect of inhibiting isocitrate dehydrogenase yields the expected accumulation of enzyme substrate (i.e isocitrate) and upstream metabolites (e.g., cis-aconitate) along with a concomitant depletion of the enzyme’s product (i.e. ketoglutarate) and downstream metabolites (e.g. succinate). In other cases, metabolite changes in response to enzyme inhibition propagate beyond reactants to distal metabolites. For instance, changes in glucose-6-phosphate and fructose-1,6-bisphosphate upon inhibition of pyruvate kinase (Supplementary fig. 3). Others exhibit transient behaviors (Supplementary fig. 2): for instance, SDH inhibition leads to only a short transient decrease of its product - fumarate levels before it comes back to its basal level - i.e. metabolite concentration before enzyme inhibition.

Independently from noise levels, we found that in some cases, enzymes sharing similar reactants, like PPC and PYK, are difficult to distinguish as perturbation targets (Supplementary Figure 3). This is evident from the fact that dynamic changes induced by inhibiting 50% of PPC or PYK activity do not lead to measurably different kinetic responses. (see Figure 1D). Similarly, simulating a partial coverage of metabolites, as would occur in experimental measurements, by removing specific metabolites from the comparison between model predictions and in silico data, reduces predictive power and makes it harder to distinguish between certain target candidates (Supplementary Figure 4-5). In our code, this can be tested by specifically masking all metabolites that cannot be measured in a given experiment (Figure 2F).

Remarkably, our simulations suggest that the qualitative behavior of the system is quite robust to changes in elasticities. Hence, the major characteristics/trends of response dynamics are mainly driven by the reference state (i.e. flux directions, reference fluxes, and metabolite concentrations, see following section), by structural network properties and by the choice of the enzyme perturbed, but much less so by the specific kinetic properties of the network. An emblematic example is the simulated inhibition of succinate dehydrogenase (SDH). IMF-simulated linear dynamics in response to partial inhibition of SDH activity are consistently more similar to the in-silico SDH inhibition dynamics than to those of any other enzyme inhibition, regardless of the sampled elasticities (i.e., across 100 different model instances) (Supplementary Figure 5).

In practice, depending on the measurement noise and on which metabolites can be measured, resolving exact enzyme targets of perturbations may be challenging. Nevertheless, in silico benchmarking showed that IMF allows us to narrow down the most likely direct targets of a perturbation despite noisy data or missing detected metabolites (Figure 1 E-F). By masking such metabolites also in artificial data, we can study specifically how this missing knowledge affects our ability to infer perturbed enzymes, or the level of certainty, thereby informing experimental design.

Accordingly, IMF may also be used for optimal experimental design, for example to determine most informative time points for measurements, accounting for the limited ability to accurately measure or distinguish certain metabolites.

### Condition-dependent response to perturbations

Microbial metabolism depends strongly on the environmental conditions, and the same enzyme inhibition, applied in different conditions, may elicit radically different metabolic changes. Different carbon sources enter at different locations in the metabolic network, leading to different flux distribution and, in some cases, different flux directions. Since enzyme inhibition typically causes upstream metabolites to accumulate and downstream metabolites to deplete (as predicted by the crossover theorem; see Introduction), we expect observable metabolic changes to depend strongly on flux directions. As a consequence, different external conditions, leading to different initial reference states and opposite flux directions, can produce qualitatively and quantitatively different metabolic responses to the *same* enzyme inhibition.

But aside from flux directions, the way perturbations propagate across the network depends also on basal metabolite concentrations and on the reaction elasticities along the way. Specifically, metabolites with high concentrations may act as buffers that block an incoming perturbation. In completely irreversible reactions - whose rates do not depend on product concentrations - perturbations cannot propagate against the flux direction, while in near-equilibrium reactions perturbations can travel relatively freely in both directions. All these effects may play out differently depending on the initial reference state. IMF takes all this into account, based on our best knowledge of the system, and uses sampling to check feasible parameter combinations or alternative hypotheses about the unperturbed metabolic state.

To test whether the model can capture metabolic responses to enzyme perturbations applied under different environmental conditions, we used fluxes and metabolic concentrations measured in *E. coli* growing on different carbon sources^67^. Based on these data, we defined 4 different reference steady states of *E. coli* growing in minimal medium with glycolytic (i.e. fructose and galactose) and gluconeogenic (acetate and pyruvate) carbon sources, giving rise to different patterns of flux directions and metabolite concentrations in central pathways (see Figure 3 and ^67^).

**Fig. 3.**
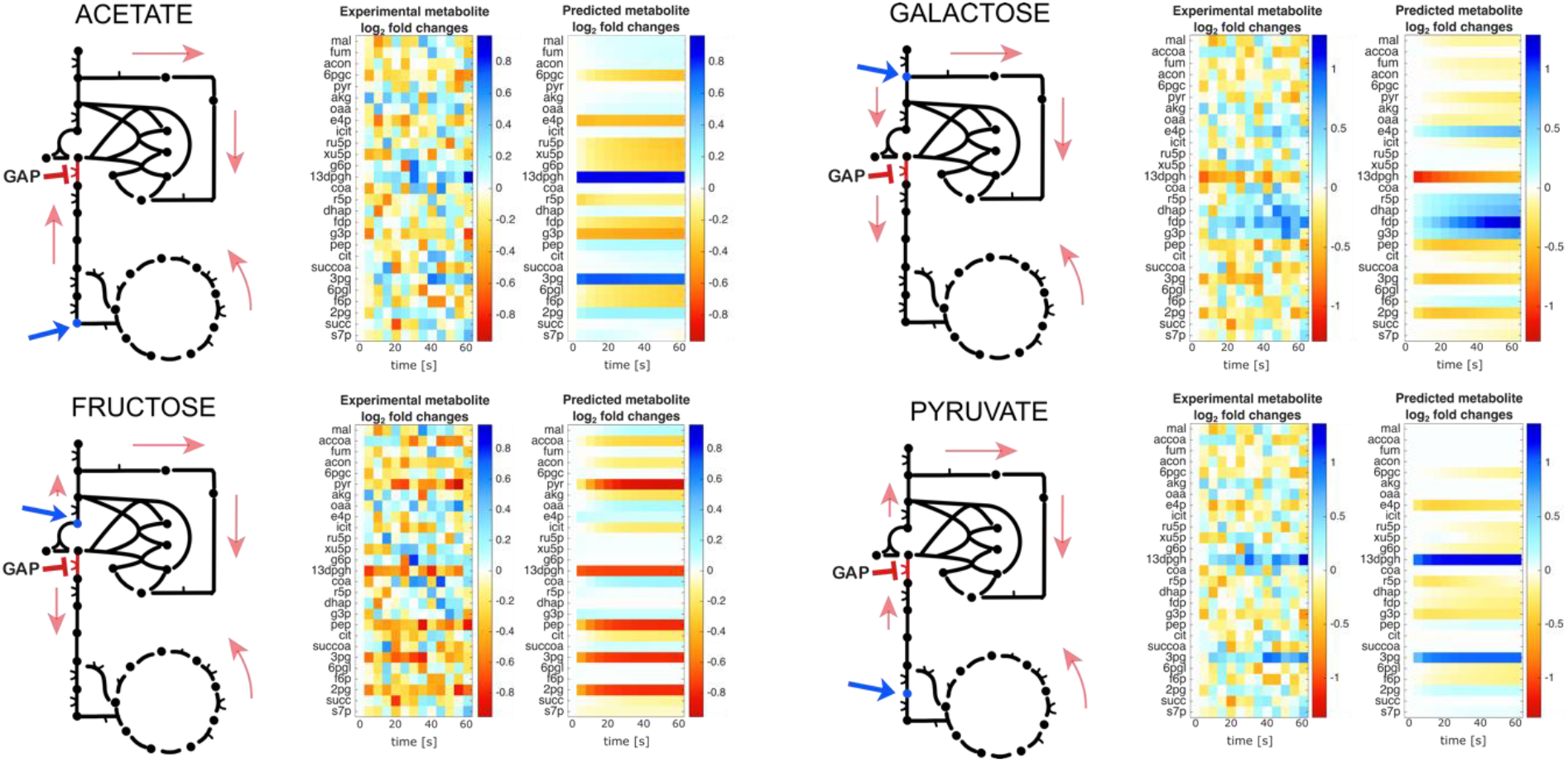
Inference or perturbed enzyme in models with different unperturbed reference states. The inhibited reaction that we chose is Glyceraldehyde 3-phosphate dehydrogenase (GAP). Depending on nutrient conditions and the resulting steady-state flux directions, the same metabolite can undergo different changes (increasing or decreasing) according to the flux direction. The blue arrow in the models indicates the point of entry of the initial metabolite in the cycle. The heatmaps display the log2 fold changes of the dynamics for all the internal metabolites (noisy in-silico data and reconstructed most likely model prediction). The horizontal direction represents the simulated time range (0, 5, 10, 15, 20, 25, 30, 35, 40, 45, 50, 55, and 60 seconds).

**Fig. 4.**
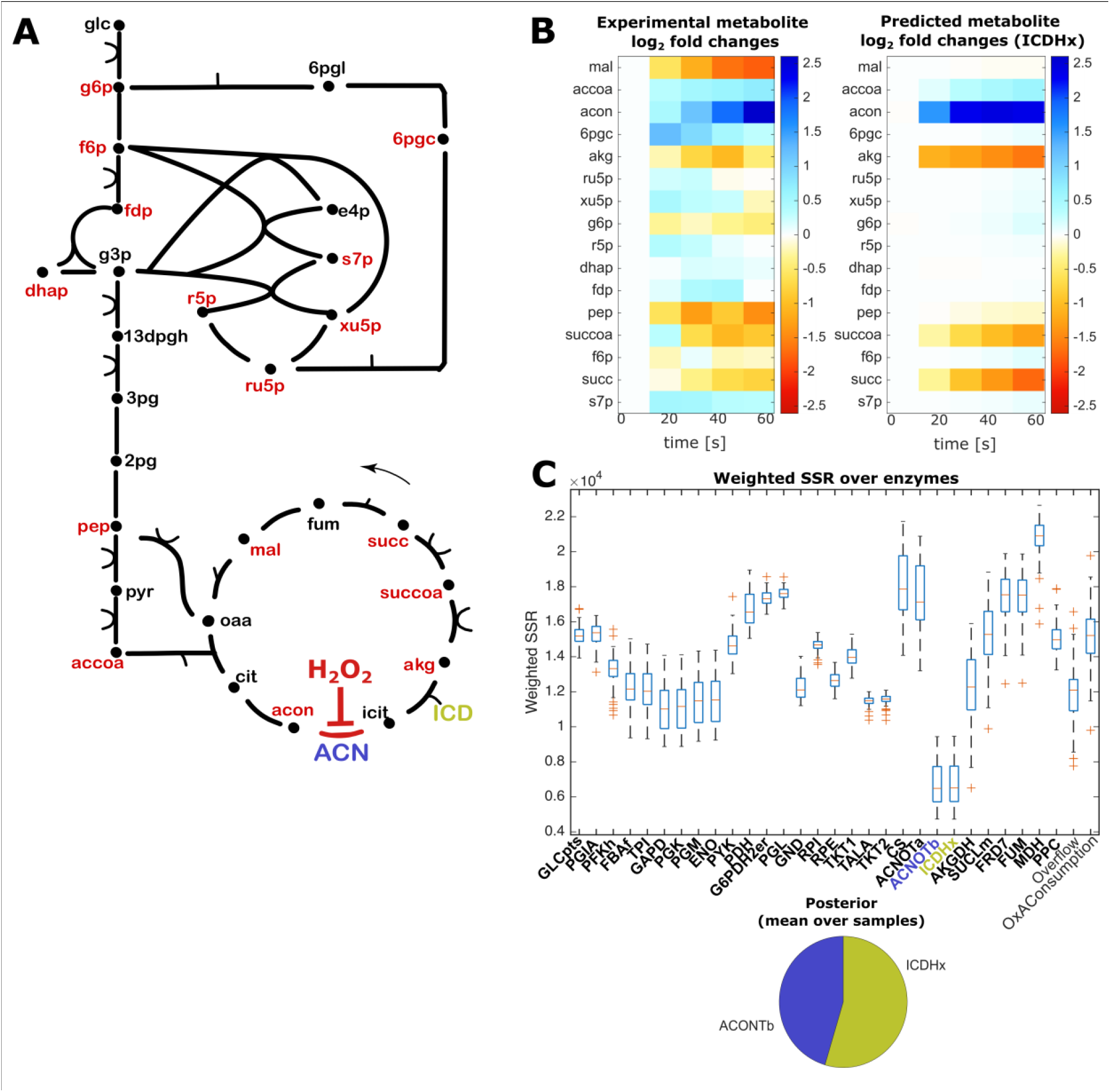
Inference of an enzyme perturbed by oxidative stress in *E. coli*. (A) Network model with measured metabolites highlighted in red. The inhibited reaction catalyzed by aconitase (ACN) and the adjacent isocitrate dehydrogenase (ICD) are marked. The black arrow indicates the flux direction in the TCA cycle under the experimental conditions. (B) Comparison between measured metabolite profiles at the onset of oxidative conditions^6^ and simulated metabolite profiles, for the maximum posterior solutions inferring the enzyme ICD as the target. The heat map shows log_2_ fold concentration changes of all the internal metabolites in a time range of 1 minute (measurements at 0, 5, 15, 30 and 60 seconds). (C) The box plot shows distributions of the negative log-likelihoods for each enzyme in the network. The pie chart shows posterior probabilities for the enzymes to be inhibited by H_2_O_2_. Only ACN and ICD have high, and in fact almost equal, probabilities.

For each nutrient condition we repeated the benchmarking approach described in the previous section for glucose minimal medium. Specifically, we (i) used the model with full kinetics to generate in silico metabolites dynamics upon 50% inhibition of each individual enzyme activity, (ii) used IMF with 100 sampled model instances and linearized metabolic dynamics to simulate the response to an inhibition of each individual enzyme, (iii) compared the model predictions to in silico data and inferred the perturbed enzyme by a maximum posterior approach based on how similar their simulated dynamics are to the original in silico data.

While the same perturbation can lead to radically different dynamic responses across different initial reference states (Figure 3), inference of the perturbed enzyme worked equally well for all the different reference states. An emblematic example is the inhibition of glyceraldehyde 3-phosphate dehydrogenase (GAP), the enzyme responsible for the reversible oxidative phosphorylation of D-glyceraldehyde-3-phosphate (G3P) to 1,3-bisphospho-D-glycerate (13GPDH). As expected, when E. coli is growing on glycolytic (e.g. galactose, fructose) carbon sources, an inhibition of GAP causes the immediate accumulation of its substrate G3P but also that of an upstream precursor in the pentose phosphate pathway (PPP): D-erythrose 4-phosphate (E4P). If instead *E. coli* is forced to grow on a gluconeogenic carbon source (e.g. acetate, pyruvate), then, as expected, we observe opposite changes: an immediate accumulation of 13GPDH and a depletion of intermediates in the PPP. It is worth noting that in all four conditions, in silico perturbations of GAP were correctly inferred by IMF.

Overall, our in-silico benchmarking results show the potential and versatility of IMF to infer the key enzyme target of a perturbation directly from dynamic metabolic data and suggest that IMF results are remarkably robust to noise in the data (Figure 2) and to different nutrient conditions (Figure 3). However, our in-silico data were still obtained using model assumptions that may not hold true in reality, thereby confounding or invalidating the simulation results (e.g. fixed concentrations of external metabolites such as cofactors, or the assumption of small perturbations). Therefore, we also tested the ability of IMF to infer the target of a perturbation from real experimental data.

### IMF analysis of *Escherichia coli*’s response to Hydrogen Peroxide

Here we applied IMF to published metabolomics data capturing absolute changes of 30 intracellular metabolites measured 5, 15, 30 and 60 seconds after challenging exponentially growing *E. coli* with 1 mM of hydrogen peroxide (H_2_O_2_), a strong oxidizer (data from ^6^).

Depending on concentration and duration, Reactive Oxygen Species (ROS) can elicit diverse effects in cells by affecting the activity of enzymes containing metal cofactors such as iron or zinc. In *E. coli*, H_2_O_2_ can oxidize and disable [4Fe-4S] dehydratases of the aconitase class, including AcnB and to a lesser extent AcnA^70^ in the tricarboxylic acid (TCA) cycle. H_2_O_2_ also targets the [4Fe-4S] isopropylmalate isomerase in leucine biosynthesis^71^, as well as mononuclear Fe(II) enzymes such as ribulose-5-phosphase 3-epimerase (RPE), an enzyme in the pentose-phosphate pathway. In both enzyme families, metal centers are accessible to direct oxidation. As a direct consequence, the dissociation of the oxidized metals inactivates the enzymes.

Although the observed reduction in α-ketoglutarate and accumulation of cis-aconitate are consistent with aconitase inhibition, several other metabolites also exhibit pronounced responses to H_2_O_2_. Nonetheless, IMF identifies aconitase (ACN) and isocitrate dehydrogenase (ICD), two adjacent enzymes in the TCA cycle next to alpha-ketoglutarate and cis-aconitate, as the two most probable direct targets of H_2_O_2_ – i.e. their inhibition yield the best fit of the experimental data.

The inability of IMF to discriminate between the two enzymes ACN or ICD as the source of the observed dynamics comes from the fact that isocitrate, the intermediate metabolite product of ACN and substrate of ICD was not measured. We had already observed the inability of IMF to uniquely identify ACN as the target of the perturbation when isocitrate was omitted from the in-silico-measured metabolites (Fig. 1F).

Remarkably, despite dynamic changes in several metabolites beyond the expected citrate accumulation and downstream α-KG depletion following H_2_O_2_ perturbation, and despite only partial experimental metabolite coverage, IMF correctly identified the primary direct target of H_2_O_2_ in central metabolism. Hence, a model-based detection of perturbed enzymes can aid data interpretation and testing of alternative hypotheses. For example, although other enzymes in central carbon metabolism (CCM), such as ribulose-5-phosphate epimerase, may be impaired by H_2_O_2_, IMF suggests that their inhibition is not the primary driver of the observed metabolic changes under the conditions tested. Nevertheless, it is also worth noting that ACN inhibition is not sufficient to explain the accumulation of intermediates in upper glycolysis like glucose 6-phosphate or fructose 6-phosphate. These changes are likely mediated by post-translational regulation of enzymes in upper glycolysis by TCA cycle intermediates, for example the inhibition of PFK by citrate. Capturing such changes would likely require to also model reactions and regulatory interactions beyond CCM^53^. While IMF can readily incorporate such interactions into model simulations, all benchmarking presented here is based solely on the core kinetic structure of the system. This choice reflects the fact that many molecular binding interactions remain only partially characterized and do not necessarily imply functional relevance.

Together, our analyses of artificial and real metabolite data show that inhibited enzymes - and in the future, possibly, drug targets - can be identified provided that metabolite data are sufficiently complete. However, different enzymes may be hard to resolve if there are no measured metabolites in between the two. In this case, different possible targets are reported together with their respective likelihoods.

## Discussion

Here we explored the analysis and interpretation of time-resolved metabolomics data by linearized kinetic models. Allowing for fast simulations, such models are well suited for computation-intense tasks such as parameter estimation or identifying perturbed enzymes. Our modeling approach describes in detail, and quantitatively, how perturbations propagate in the network.

Applying IMF to the central metabolism of *E. coli* as a test case, we showed that linearized kinetic models can accurately represent the system-wide, short-term response to a sudden perturbation and, as a result, can help us infer the origin of the perturbation. The models are built semi-automatically from a given network structure, estimated unperturbed metabolic states, and a sampling of kinetic constants via reaction elasticities and saturation values.

To construct the unperturbed steady state, IMF relies on known metabolic fluxes and uses the flux directions to obtain thermodynamically feasible metabolite concentrations close to the data. Since fluxomics data may be hard to obtain, estimates of intracellular fluxes may be determined from a flux balance model constrained by exometabolomics data of nutrient uptake rates and byproduct secretion rates^72–75^. After determining the unperturbed reference state, IMF simulates how perturbations caused by an impacted enzyme would spread in the network.

A key assumption of our approach is that protein levels remain unchanged. This eliminates a major source of uncertainty from the model, but it also restricts the analysis to short-term metabolic responses. Mathematically, both short-term and long-term responses are described by the time-dependent control coefficients, which underlie our simulations. For very short times t ≈ 0, the control matrix **C**^*s*^(t) can be approximated by **C**^*s*^(t) ≈ t **N** (a sparse matrix indicating that only direct substrates and products of the perturbed reaction are affected); in contrast, for long times t → ∞, the formula converges to the control matrix **C**^*s*^ from MCA, which describes changes of steady-state concentrations in the entire system.

A limitation of IMF lies in the assumption of small deviations from the initial steady state, which means that only mild inhibitions can be applied. This assumption might not hold in experimental scenarios with perturbations such as gene knockdowns (e.g. CRISPRi or RNAi) or high-dosage drug treatments. While this limitation could be overcome by using full kinetic models, the numerical effort of such models is larger and scales poorly with the size of the model. Remarkably, our in-silico simulations showed that even relatively strong inhibitions of enzyme activity are well captured, at least qualitatively, by a linearized model. It is also worth noting that enzyme inhibitors do not act instantaneously, but must first enter the cell and engage their target. Similarly, following transcriptional inhibition, degradation of the gene product and dilution through cell growth lead to a progressive reduction in its activity. In our model this might be taken into account by describing the drug as a metabolite that is transported into the cell and inhibits the reaction catalyzed by its enzyme target, or by an inhibitory function that scales with the doubling time or degradation rates of proteins.

Another limitation of IMF, as described here, is the assumption that only a single enzyme activity is being perturbed while all others remain unchanged. In reality, a perturbing agent such as a drug or a toxic compound may have multiple targets. In the model, this means that several enzymes would be perturbed simultaneously and to different extents. While this is easy to handle in forward simulations - especially in the linear approximation, where the effects on different enzymes can be obtained by a simple linear superimposition of individual enzyme perturbations - in practice inferring several perturbed enzymes is much more difficult. The difficulty arises not only from combinatorial complexity (which is largely mitigated by the linear regime), but more importantly from the difficulty to reliably distinguish between equally probable solutions that produce similar metabolic responses.

Just like MCA, in the 1970s, replaced the crossover theorem by a general quantitative theory, our MCA-based method replaces rules of thumb for interpreting metabolomics data by a quantitative modeling approach. Instead of looking for a single typical signature (metabolite accumulation or depletion around one enzyme), we make full use of data by considering all metabolites at the same time in the context of a realistic dynamical model. By taking into account metabolites from the entire network, IMF can provide strong evidence for a certain hypothesis and rule out others. Even if none of the simulated enzyme inhibitions matches the data perfectly well, by comparing evidence for and against different enzymes and by accounting for the way in which perturbations might spread, IMF is able to make good predictions even from incomplete and uncertain data. Furthermore, if no clear conclusions can be made - as in the case of the oxidative stress data - alternative explanations are provided and the remaining uncertainty can be quantified.

Although our results suggest that the qualitative behavior of the system can be relatively robust to the specific choice of elasticities, this should not be interpreted as eliminating the need for elasticity sampling. For example, setting all saturation values to 0.5 may provide a useful first approximation or starting point, and may even yield realistic predictions in some cases; however, such a fixed choice is not guaranteed to satisfy dynamical stability criteria and may lead to unstable model instances. This issue is expected to become increasingly important for larger models, where identifying combinations of kinetic parameters and elasticities that yield stable reference states becomes more delicate. In addition, sampling elasticities allows the model ensemble to explore a broader range of plausible dynamic behaviors, thereby making IMF more generalizable and robust when applied to larger and less well-characterized metabolic networks.

This ability to sample, filter, and compare alternative elasticity sets also suggests a further use of IMF beyond target inference. Another potential application of IMF is the prediction of *in vivo* kinetic parameters (e.g. Michaelis-Menten values). After probing the system with a perturbation, the resulting most likely elasticities can be converted back into conventional kinetic constants. With the increasing availability of large-scale metabolomics datasets profiling the effects of multiple perturbations, like chemical (e.g. drugs) ^8,9^ or genetic perturbations (e.g. genetic inactivation) ^76^, IMF might be used to find the set of elasticities that provides the best consensus fit across all the perturbations simultaneously. It is also worth noting that our approach can be easily extended to compartmentalized eukaryotic models, including potentially models of human cells. Some metabolites can only be measured as a sum, and IMF could be easily extended to incorporate such measurements by modifying the metric to assess similarity between measured and inferred metabolic dynamics.

Finally, a key advantage of IMF is that the accuracy of enzyme identification from metabolite data can be assessed by simulations with artificial data and inform the experimental design. This may help us anticipate if a given enzyme can be identified at all as a target of a perturbation, given the accuracy and completeness of metabolomics data; and also, what additional data would have to be collected to reduce the remaining uncertainty.

## Methods

This section gives a short description of the IMF framework including model construction, simulation, and the inference of perturbed enzymes. A more detailed description can be found in the documentation at our GitLab repository (see Code Availability).

### Modeling framework

To simulate dynamic metabolic responses to a sudden change in an enzyme activity, we use kinetic metabolic models. Following the SKM framework^**48**^, we predefine reference fluxes and metabolite concentrations, representing an unperturbed steady state, and specify a linearized dynamics around this state. A model is then parameterized by reaction elasticities, defined as derivatives of the enzymatic rate laws. In general, their values depend on various factors including the enzymatic rate laws, the kinetic constants in these laws, and the metabolic reference state considered. In our SKM models, following^**50**,**64**^, the elasticities are parameterized by unitless saturation values which can be chosen independently in the range [0,1]. By drawing these numbers from random distributions, we generate an ensemble of model instances with a predefined reference state (or alternatively, the reference states can be sampled as well). The saturation values (possibly together with the thermodynamic forces known from the reference state, if thermodynamically consistent rate laws are desired), define physically valid models which realize the reference state in question. Since their kinetic constants can be easily reconstructed, all our calculations can either be run with the linearized kinetics (directly based on reaction elasticities) or with full nonlinear rate laws (and kinetic constants derived from the same elasticities). In practice, when the number of simulations is large - e.g. in estimation or inference tasks - we use the linearized model because of its much lower numerical effort.

### Model definition and construction

A model instance is constructed from information about network structure, about the steady reference state, and about the reaction elasticities in this state.

1. The network structure is defined by a list of enzyme-catalyzed reactions (described by a stoichiometric matrix), a choice of metabolites treated as external (i.e. with constant concentrations), and a matrix defining the locations and signs of regulation arrows.
2. The reference state is described by a metabolic flux distribution and a vector of metabolite concentrations. If thermodynamic correctness is desired, the equilibrium constants (or, equivalently, standard Gibbs free energies of reaction) must be given, and the resulting thermodynamic driving forces (determined from equilibrium constants and metabolite concentrations) must match the flux directions.
3. The unscaled reaction elasticities can be obtained from the reference state (fluxes and concentrations) and the scaled reaction elasticities. The scaled elasticities, in turn, can be parameterized by saturation values, describing to what extent an enzyme is saturated with bound substrates or products. Depending on the purpose of modeling, these saturation values can be predefined or sampled. The formulae for scaled elasticities as functions of saturation values depend on the type of rate laws assumed ^50,64^. If thermodynamically consistent, reversible rate laws are used, the scaled elasticities depend on the saturation values and, additionally, on the thermodynamic driving forces. Enzyme regulation (e.g. allosteric regulation or competitive inhibition) can be taken into account: scaled elasticities for regulators are computed from the regulation matrix and a matrix of saturation values for the regulation arrows.
4. For reactions with many substrates or products (e.g. the biomass reaction) that are not usual enzymatic reactions, we rescale the sampled scaled elasticities to a sum of absolute values of 1. In this way, we avoid that the reaction becomes ultrasensitive to a general increase in metabolite concentrations, which might lead to unstable models.
5. A model instance, computed in this way, allows us to compute (1) the reconstructed enzyme kinetic constants, and therefore the full enzymatic rate laws; (2) the Jacobian matrix, and (3) the response and control matrices
6. Based on a network structure and a given reference state, model instances can be created by sampling the saturation values, then computing the elasticities (assuming a certain type of rate laws), and deriving from them all dynamic properties of the model instance (kinetic constants; Jacobian matrix; stability; and static and time-dependent control and response coefficients). By sampling many model instances, a model ensemble can be created.
7. Along with the elasticities, we can also sample the reference states. In this case, we first consider a “standard” reference state with feasible fluxes and metabolite concentrations. Then we draw a tentative flux for each reaction and a tentative concentration for each metabolite from independent log-normal distributions centered around their standard values. Finally, we apply a correction procedure to convert them into a new, consistent steady state with stationary fluxes and with metabolite concentrations leading to feasible driving forces.

In our sampling procedure, all model instances with unstable reference states are discarded. If a reference state is unstable, this means that upon certain small perturbations of metabolite concentrations, the system’s own dynamics drives the system away from the reference state. Mathematically, this happens when the Jacobian matrix has eigenvalues with real parts larger than the assumed cell growth rate *λ*(with a default value of *λ*= 1/*h*); the growth rate enters the formula because we assume dilution in growing cells, which has a slightly stabilizing effect. Conversely, a dynamical system is asymptotically stable if all Jacobian eigenvalues have real parts smaller than $\lambda$. The stability of a model instance depends on its reaction elasticities. For a given model structure, it can happen that the model instance with all saturation values set to 0.5 is unstable, but a substantial number of other (sampled) model instances may still be stable.

For further details see the documentation at our GitLab repository (see Code Availability).

### Model simulations

The inference of perturbed enzymes from metabolic fingerprints relies on simulations of the rapid metabolite dynamics after a sudden decrease in an enzyme activity. To model an enzyme inhibition, we consider the “enzyme level” *e*, which describes the enzyme activity under the effect of the drug, while other regulatory effects - by metabolites described by the model - are included in the rate laws. After a drug is applied at time *t*=0, the enzyme level suddenly switches from its initial value to a lower constant value. Starting from an unperturbed enzyme level e, the inhibition leads to a (negative) perturbation *δe*∈ [−*e*, 0], which is described by the inhibition factor 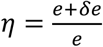. In the linear approximation, the inhibition results in metabolite log-fold changes 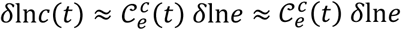 with a time-dependent control coefficient matrix 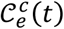. The simulation, based on these matrices, yields a metabolic fingerprint matrix containing log-fold changes of the metabolite concentrations with respect to the initial state, for a number of predefined time points and a set of “measurable” metabolites, that is, either all internal metabolites in the model or a selection of metabolites measurable in experiments.

For a model with full kinetics, the dynamic equations based on the rate laws have to be integrated numerically. While this is possible with our code, the calculations may be slow. The faster alternative is based on the linearized version of the model, which assumes small perturbations. In this case, an enzyme perturbation can be directly mapped to the resulting metabolite dynamics via scaled, time-dependent response coefficients (or control coefficients), which refer to log concentration data. The linearized differential equation system can be directly integrated by using the matrix exponential function, which is numerically much more efficient and can be directly evaluated for a desired time point without having to integrate the system along the time interval before.

### Inferring enzyme inhibitions from metabolic fingerprints

We aim to infer which enzyme in a metabolic network has been perturbed based on time-resolved metabolite measurements. An enzyme perturbation induces transient changes in metabolite concentrations, which propagate through the network and create a characteristic dynamic response. We refer to this response as the *metabolic fingerprint* of the perturbation. To infer an inhibited enzyme from its resulting metabolic fingerprint, we simulate an inhibition of each enzyme in the model and compute the corresponding predicted fingerprints. These predicted fingerprints are then compared to the experimentally observed fingerprint, and the enzyme whose prediction best matches the data is selected as the most plausible perturbed enzyme. While this idea is conceptually straightforward, several complications must be addressed in practice. First, model parameters such as reference states and reaction elasticities are typically uncertain, implying that predicted fingerprints will vary across plausible model realizations. Second, the notion of “best match” between fingerprints requires a principled definition that accounts for measurement noise and model uncertainty. Third, the strength of the perturbation is generally unknown and must be estimated jointly with the enzyme identity. To address these challenges, we formulate the inference problem in a likelihood-based framework and approximate Bayesian inference using an ensemble of sampled model instances. In summary, the inference procedure consists of the following steps:

1. Sample an ensemble of model instances reflecting uncertainty in parameters (elasticities and possibly reference states).
2. For each enzyme perturbation and each model instance, compute the predicted metabolic fingerprint.
3. Estimate the inhibition strength and compute the weighted residuals between predicted and observed fingerprints.
4. Convert residuals into likelihood values.
5. Aggregate likelihoods across model instances to approximate posterior probabilities.
6. Rank enzymes according to their posterior probabilities to identify the most plausible perturbed enzyme(s).

### Accounting for model uncertainty

A central challenge in the inference problem is that the model parameters are not precisely known. In particular, reaction elasticities and, in some cases, reference metabolite concentrations may be uncertain or entirely unknown. As a consequence, each plausible parameter set leads to a different predicted fingerprint for the same enzyme perturbation. To account for this uncertainty, we generate an ensemble of model instances ℳ_***s***_ (***s*** = **1**, … , ***N***) by sampling parameter values from suitable distributions. For example, if no prior knowledge about elasticities is available, they may be drawn independently from uniform distributions over biologically plausible ranges. Similarly, reference states may be sampled around a predefined baseline state to reflect uncertainty. Each sampled model instance yields a different predicted fingerprint for each enzyme perturbation. The inference procedure therefore compares the observed fingerprint not to a single prediction, but to a distribution of predictions across model instances.

### Metabolic fingerprints

Consider a metabolic network with metabolite concentrations ***c***_***i***_(***t***) and enzymes indexed by ***r***. A perturbation of enzyme ***r***, characterized by an inhibition factor ***η*** ∈ (**0, 1**), induces time-dependent changes in metabolite concentrations. We define the metabolic fingerprint as the matrix of log fold changes

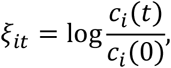

measured for metabolites *i* at time points *t* > 0. The initial time point is omitted since ξ_*i*0_ = 0 by definition. Given a model instance ℳ_*s*_, defined by a reference state and kinetic parameters (e.g. elasticities), we simulate the system’s response to a perturbation of enzyme *r* and obtain a predicted fingerprint 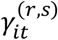. For small perturbations and under a linear approximation on logarithmic scale, fingerprint amplitudes scale approximately with the perturbation strength,

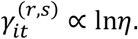

### Comparison of predicted and observed fingerprints

Each fingerprint, originally defined as a matrix over metabolites and time points, is represented as a vector by concatenating its entries. This allows the use of standard distance and similarity measures. To quantify the discrepancy between predicted and observed fingerprints, we consider the weighted sum of squared residuals (wSSR),

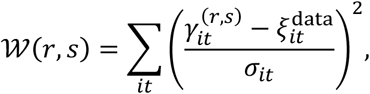

where *σ*_*it*_ denotes the standard deviation of measurement noise. This measure corresponds to a weighted Euclidean distance and ensures that data points with smaller uncertainty contribute more strongly to the comparison.

### Likelihood interpretation

Rather than treating this comparison as an ad hoc distance measure, we interpret it statistically. Assuming independent Gaussian measurement noise,

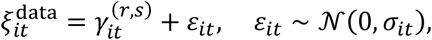

the likelihood of enzyme *r* for model instance *s* is

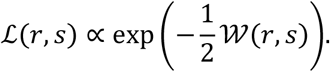

Thus, maximizing the likelihood is equivalent to minimizing the weighted residuals. This formulation provides a principled way to compare fingerprints while accounting for measurement uncertainty.

### Estimation of inhibition strength

The inhibition strength ***η*** is generally unknown and must be estimated for each enzyme and model instance. There are two alternative methods. The first method is based on a Maximum-likelihood estimation. Assuming approximate linear scaling of fingerprints with **ln*η***, the optimal inhibition strength is obtained by minimizing 𝒲, yielding

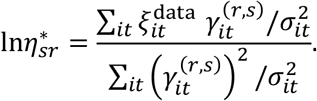

This estimator corresponds to a weighted linear regression of the observed fingerprint onto the predicted fingerprint. The second, approximate method is based on data normalization: predicted and observed fingerprints are normalized by their Euclidean norms prior to comparison. This removes differences in amplitude and focuses on matching the shape of the fingerprints. The inhibition strength can then be approximated by rescaling the simulated fingerprint to match the norm of the data. This approach is exact under linear scaling assumptions and remains effective as a heuristic in more general settings.

### Posterior formulation

The inference problem can be formulated in a Bayesian framework. Let ℳ denote the model parameters. The posterior distribution over enzyme indices is obtained by marginalizing over model uncertainty,

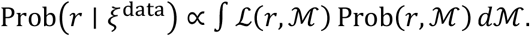

Assuming a uniform prior over enzymes (i.e. all enzymes are a priori equally likely targets) and a prior over model parameters consistent with the sampling procedure, this simplifies to

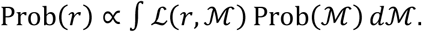

### Approximation by sampling

In practice, we approximate this integral using the ensemble of sampled model instances,

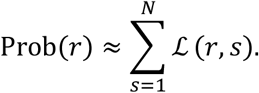

This corresponds to averaging likelihood contributions across all plausible models. As a computationally simpler alternative, one may also approximate the integral by selecting the maximum likelihood across samples. The resulting values are normalized across all enzymes to obtain posterior probabilities.

### Choice of most plausible perturbed enzyme or enzymes

The posterior probability of an enzyme reflects how well its predicted fingerprints explain the data, averaged over model uncertainty. By ranking enzymes according to these probabilities, we obtain a list of candidate targets. Based on the posterior probabilities, one may identify either a single most plausible enzyme or a set of enzymes whose cumulative probability exceeds a chosen credibility threshold.

### Dependence of posterior probabilities on measurement uncertainty

An important aspect of the inference is its dependence on the assumed measurement uncertainties ***σ***_***it***_. These values directly affect the likelihood and therefore the posterior probabilities. If all error bars are increased, the likelihood differences between enzymes become smaller, leading to more uniform posterior distributions. Conversely, smaller error bars result in more peaked posteriors, concentrating probability on fewer enzymes. Importantly, while the absolute posterior values depend on the error scale, the ranking of enzymes is typically preserved. This highlights the importance of choosing realistic error estimates when interpreting posterior probabilities.

### Numerical effort

For the results presented here, we performed all the computations on an iMac 2017 equipped with a 4.2 GHz Quad-Core Intel Core i7 processor, 16 GB of 2400 MHz DDR4 RAM, running macOS 13.7. The inference procedure using a hundred different model instances required approximately 7 seconds.

### Tests with artificial data

To test IMF with artificial data (“in silico data”) we run simulations for a “true” model with a given reference state and elasticity values. To simulate measurement noise, independent Gaussian-distributed noise is added to the data values. When generating artificial data, different choices can be made: (1) Simulation using a “true model” full or linearized kinetics; (2) different experimental noise levels; (3) masking of “unmeasurable” metabolites, e.g. the metabolites that are not measured in the experimental data set of interest. By generating artificial data from nonlinear models, assuming higher noise levels, and masking metabolites, users can make the estimation problem increasingly realistic, but also increasingly hard. In the results shown above, we used a model with nonlinear rate laws, a realistic reference state, kinetic constants reconstructed from predefined saturation values, and realistic experimental noise levels (with a geometric standard deviation of 1.1 for individual metabolite measurements, corresponding to about 10% measurement error).

In tests with in-silico data, the model used for inference may be identical to or different from the “true” model for generating the data. In the first case, we imply that we have full knowledge of the true metabolic model - or are lucky enough to guess it right - while in the second case we assume, realistically, that our model used for inference does not faithfully describe reality.

### Models and data

The models and data used in this work are available at our GitLab repository (see Code Availability).

### *Escherichia coli* model and data

To model *E. coli* core metabolism, we used the model from ^65^ in a modified version. The modifications concern a few draining reactions which mimic side branches and enable the model to describe measured fluxes while assuming a steady state. A steady state was not required in^65^ , but it is required here for a meaningful description of enzyme perturbations. A standard reference state for aerobic growth on glucose was constructed from fluxomics data ^66^ and metabolomics data ^67^, mapped to the model as in ^65^. Metabolic reference states for growth on other carbon sources were constructed based on data from ^67^.

## Further information

### Code availability

Matlab code for IMF (developed and tested on Matlab 2019b) as well as example models and data are available at our GitLab repository https://git.scicore.unibas.ch/zampieri-lab/inference-from-metabolic-fingerprints under a GNU Public License.

## Acknowledgements

We are thankful for funding by SNF Sinergia (CRSII5_189952), NCCR AntiResist project funding (180541), Novartis Forschungsstiftung (FN24-0000000612), NIH Research Project (R01) (1R01AI173328-01), the German Research Foundation (LI 1676/2-1), the Agence Nationale de la Recherche (ANR-21-CE45-0021-02), and the PEPR B-BEST (project MuSiHC). We gratefully acknowledge the support of sciCORE scientific computing core facility at the University of Basel and the Euler cluster operated by the High Performance Computing group at ETH Zürich. We thank both facilities for providing the computational resources and support.

## Contributions

WL + MZ: developed method; CR: contributed to implementation and ran model simulations; WL + MP: implemented method; MP + TD ran simulations and generated figures; all authors contributed to preparing the manuscript.

