## Supplementary Figures for "Model-based inference of enzyme inhibitions from perturbation-induced metabolic dynamics"

Wolfram Liebermeister[1+*], Michela Pauletti[2,3 ∘+], Terézia Dorčáková[2], Caspar Rahm[3], and Mattia Zampieri[2,3*]

[1]Université Paris-Saclay, INRAE, MaIAGE, 78350 Jouy-en-Josas, France
[2]Department of Biomedicine, University of Basel, Basel, Switzerland
[3]Institute of Molecular Systems Biology ETH Zürich, Zürich, Switzerland

[+] Contributed equally to this work

[∘] Current address: Interventional and Experimental Endoscopy (InExEn), Department of Internal Medicine 2, Universitätsklinikum Würzburg, Germany

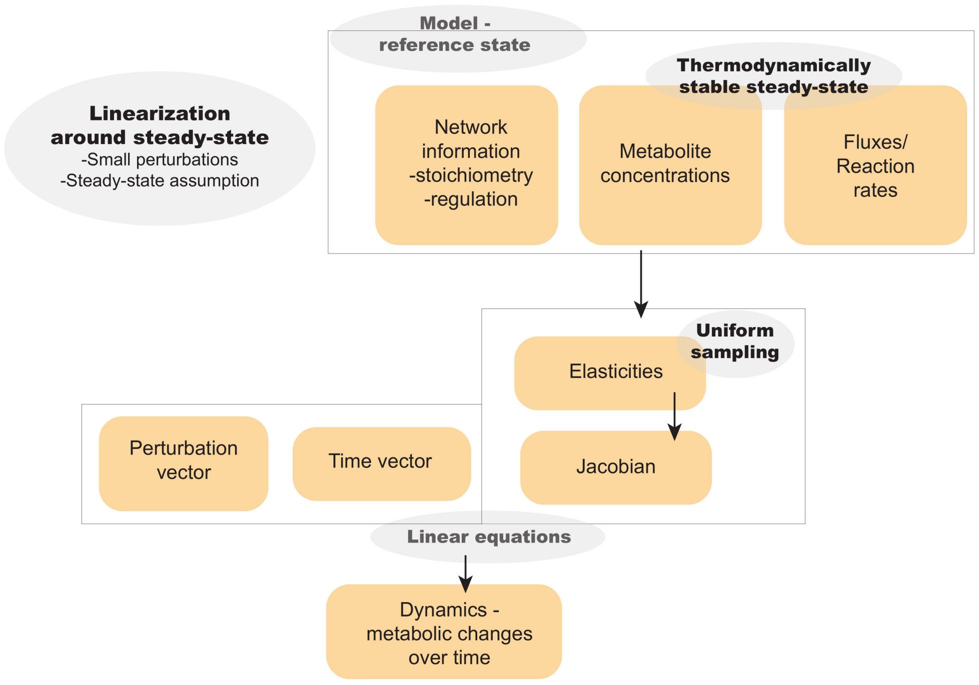


Figure S1: Structural Kinetic Modeling (SKM) flowchart - The diagram shows the ingredients, some assumptions, and the steps required by SKM as used in our Inference from Metabolic Fingerprints (IMF) method.

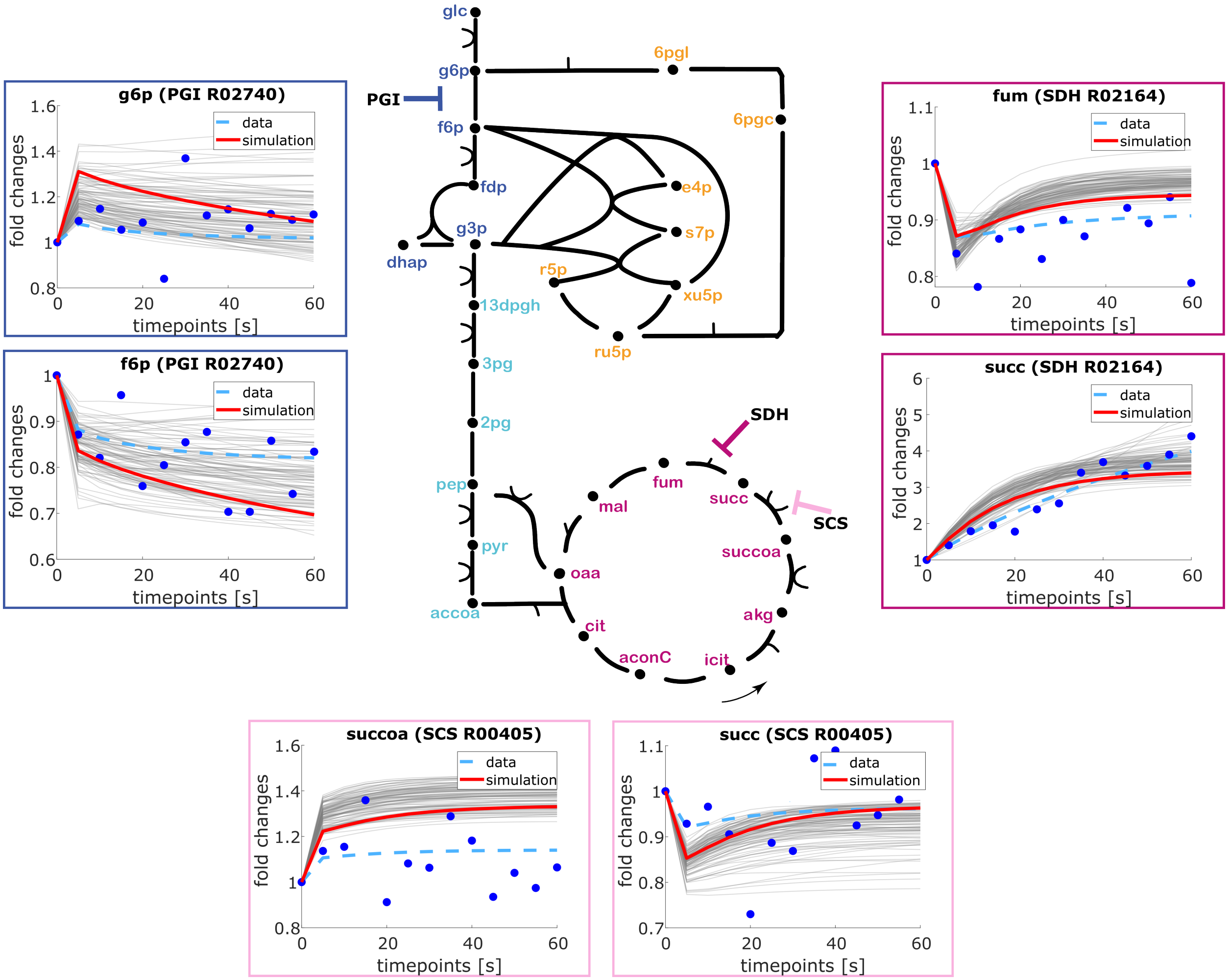


Figure S2: Simulation of enzyme perturbation leads to distinct substrate and product dynamics - Using the same set up as described in Figure 1, we run SKM to simulate the metabolite dynamics of 50% inhibition of the PGI (glucose-6-phosphate isomerase), SDH (succinate dehydrogenase) and SCS (succinyl coenzyme A synthetase) catalyzed reaction. In the graphs, the grey lines represent the metabolite dynamics in all simulations of a specific inhibition, the red line highlights the simulation which fits the target data best when all the metabolite dynamics are considered. The light blue dashed line represents the target data and the blue dots indicate the target data dynamics after adding 10% noise. While the inhibition can lead to persistent substrate accumulation (succinate - succ in SDH inhibition and succinyl-CoA - succoa in SCS inhibition) or product depletion (fructose-6-phosphate - f6p in PGI inhibition), we also observed transient changes in metabolite levels such as short-term accumulation of glucose-6-phosphate - g6p when PGI is inhibited or temporary decrease of succ levels upon SCS inhibition and fumarate - fum upon SDH inhibition.

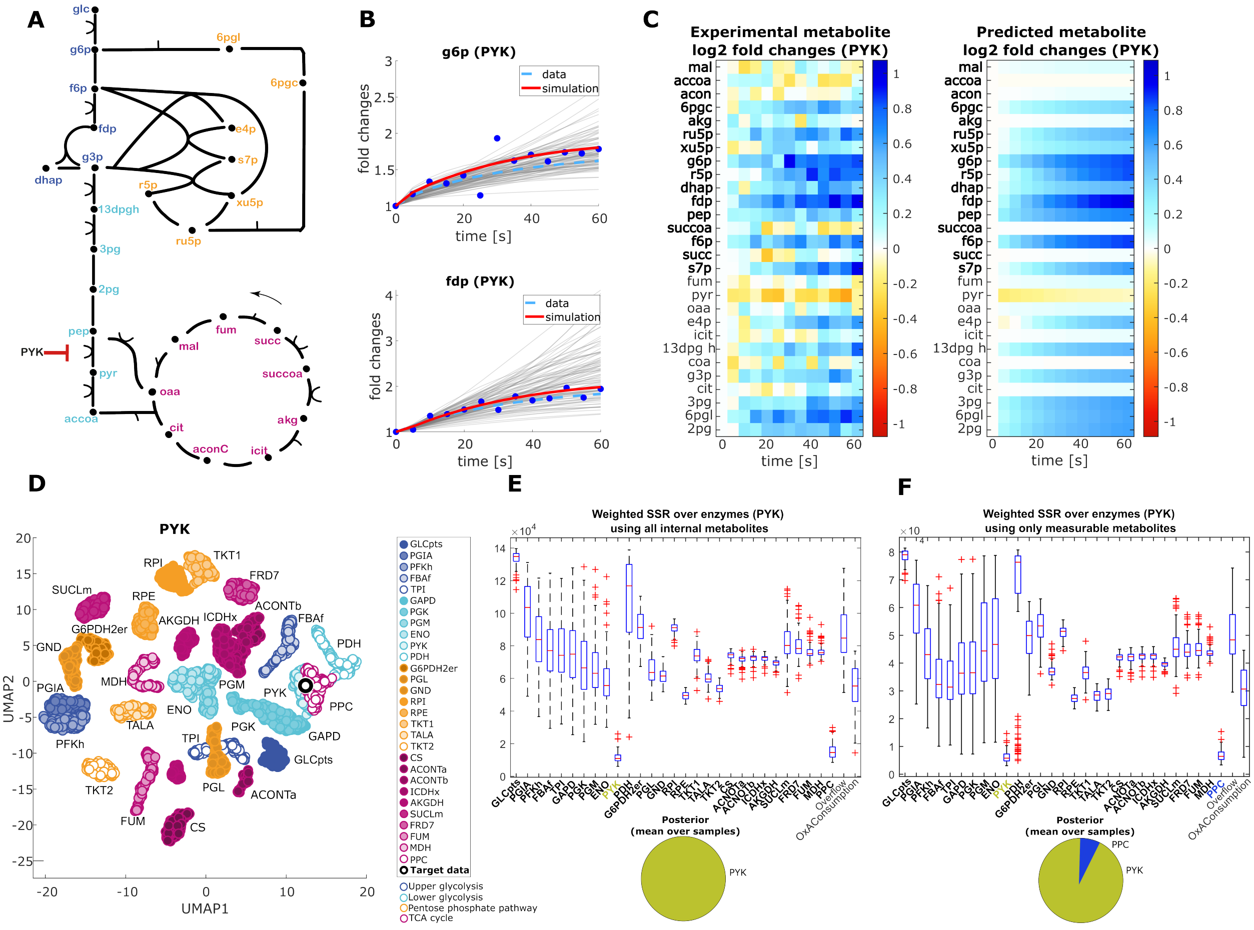

Figure S3: Enzyme perturbation leads to unexpected changes in the levels of distant metabolites - Using SKM with the same configuration as described in Fig.1, we simulated 50% inhibition of PYK (pyruvate kinase). A - The E. coli model of metabolism used to generate the data shown in this figure contains 31 reactions and 40 metabolites (out of which 12 are considered external, showing fixed concentrations). B - The dynamics of metabolites distant to the inhibited reaction (glucose-6-phosphate - g6p and fructose-1,6-bisphosphate - fdp). The grey lines represent the g6p/fdp dynamics in all simulations of PYK inhibition, the red line highlights the simulation which fits the target data best when all the metabolite dynamics are considered, the light blue dashed line represents the target data and the blue dots indicate the target data dynamics after adding noise. C - The heatmaps show the dynamics of all the internal metabolites of the target dataset (left) and the closest simulation (right). D - We projected the metabolite dynamics of all the hundred simulations of inhibition of each reaction in the network using UMAP. In the same space, we calculated the embeddings of the target data (black edge). E, F - The box plots show the distributions of the noise-weighted distances of the hundred simulated dynamics of every reaction in the network to the target data calculated using either all the internal metabolites (E) or only the measurable metabolites (measurability determined by the dataset from figure 6, highlighted in the heatmaps (C) in bold) (F). The pie charts show the posterior probability of a reaction being inhibited.

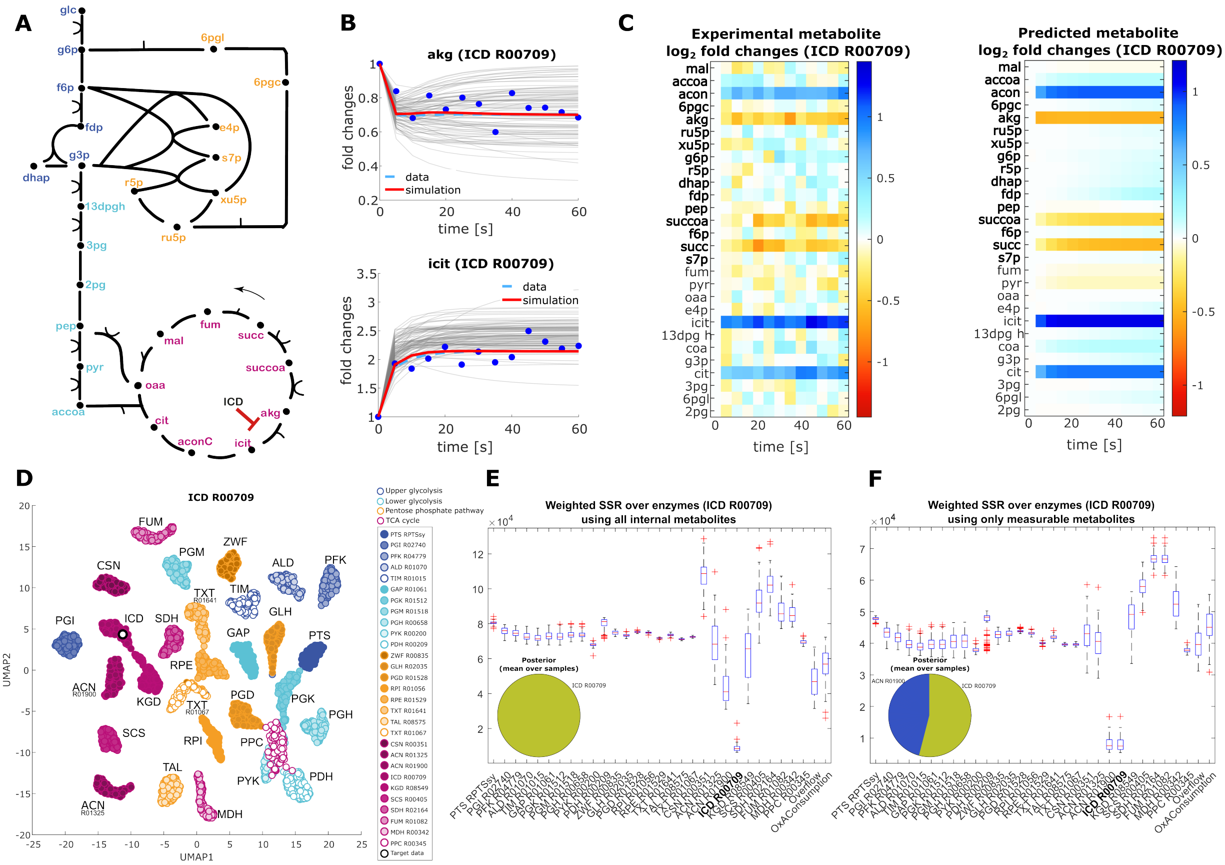

Figure S4: Simulated perturbation of the enzyme ICD and inference of perturbed enzyme from artificial data - See the corresponding Figure 1 (for a perturbation of ACN) in the main article.

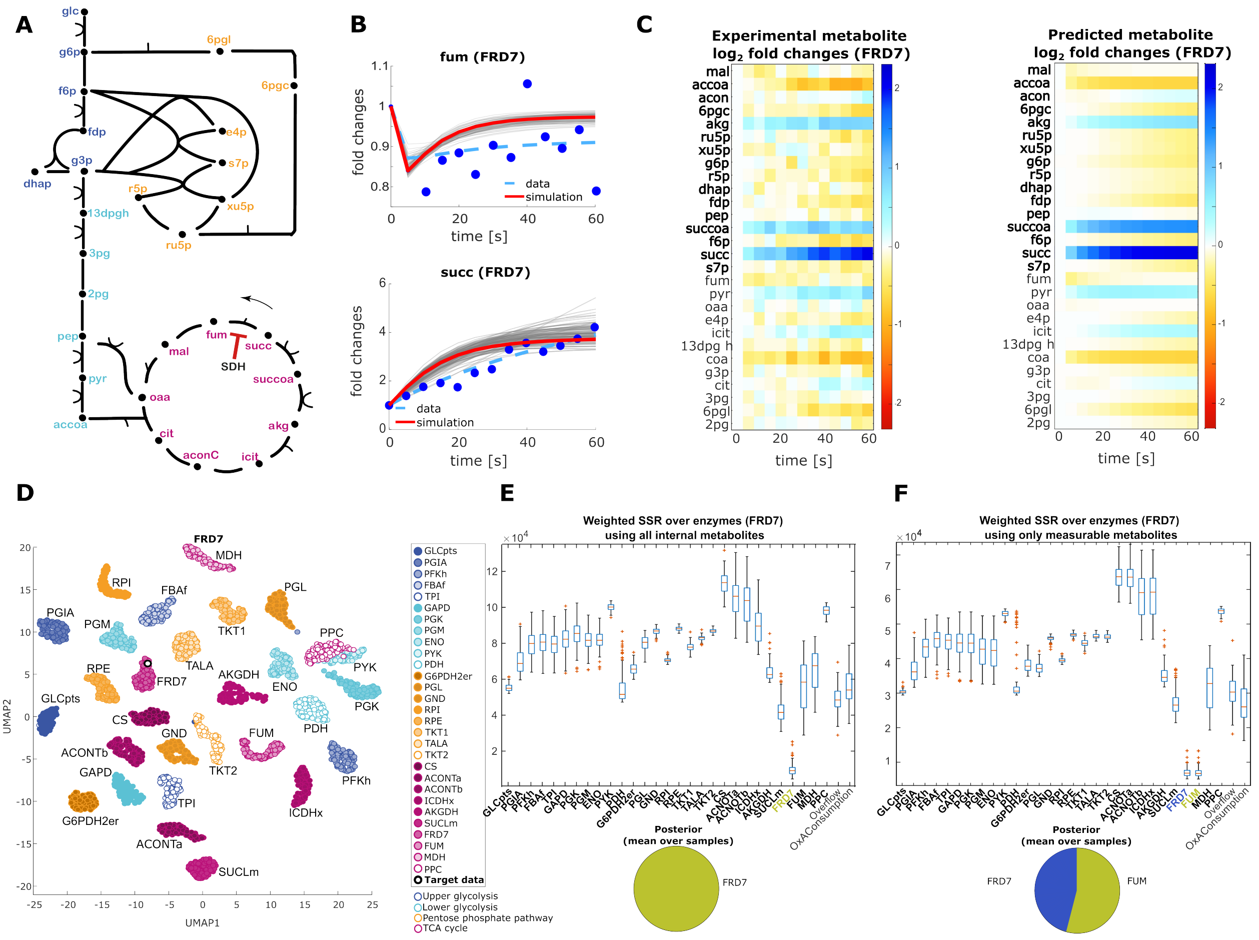

Figure S5: Simulated perturbation of the enzyme SDH and inference of perturbed enzyme from artificial data - See the corresponding Figure 1 (for a perturbation of ACN) in the main article.
